# CRISPR interference for Sequence-Specific Regulation of Fibroblast Growth Factor Receptor A in *Schistosoma mansoni*

**DOI:** 10.1101/2022.08.17.504262

**Authors:** Xiaofeng Du, Donald P. McManus, Juliet D. French, Natasha Collinson, Haran Sivakumaran, Skye R. MacGregor, Conor E. Fogarty, Malcolm K. Jones, Hong You

## Abstract

Employing the flatworm parasite *Schistosoma mansoni* as a model, we report the first application of CRISPR interference (CRISPRi) in parasitic helminths for loss-of-function targeting the *SmfgfrA* gene which encodes the stem cell marker, fibroblast growth factor receptor A. SmFGFRA is essential for maintaining schistosome stem cells and critical in the schistosome-host interplay. The *SmfgfrA* gene was targeted in *S. mansoni* adult worms, eggs and schistosomula using a catalytically dead Cas9 (dCas9) fused to a transcriptional repressor KRAB. We showed that *SmfgfrA* repression resulted in considerable phenotypic differences in the modulated parasites compared with controls, including reduced levels of *SmfgfrA* transcription and decreased protein expression of SmFGFRA, a decline in EdU (thymidine analog 5-ethynyl-2’-deoxyuridine, which specifically stains schistosome stem cells) signal, and an increase in cell apoptosis. Notably, reduced *SmfgfrA* transcription was evident in miracidia hatched from *SmfgfrA*-repressed eggs, and resulted in a significant change in miracidial behavior, indicative of a durable repression effect caused by CRISPRi. Intravenous injection of mice with *SmfgfrA*-repressed eggs resulted in granulomas that were markedly reduced in size and a decline in the level of serum IgE, emphasizing the importance of SmFGFRA in regulating the host immune response induced during schistosome infection. Our findings show the feasibility of applying CRISPRi for effective, targeted transcriptional repression in schistosomes, and provide the basis for employing CRISPRi to selectively perturb gene expression in parasitic helminths on a genome-wide scale.

## 1 Introduction

*Schistosoma mansoni* is a flatworm parasite that causes schistosomiasis, a disease which afflicts 250 million people in 74 countries [1–4].Currently, no anti-schistosome vaccines are available for human use and clinical treatment relies entirely on the single drug praziquantel (PZQ). The potential emergence of PZQ drug resistance is an ever-present concern [1]. Effective vaccines and new treatments for reducing the global burden of schistosomiasis are thus needed urgently. The past few decades have witnessed new advances in schistosome developmental biology, genomics, proteomics and transcriptomics, and in our understanding of schistosome-induced pathogenesis, and the host-parasite interaction [1, 2, 5–10]. Notably, complete genomic sequences of the three main schistosome species (*S. mansoni* [5, 6], *S. japonicum* [7, 8] and *S. haematobium* [9, 10]) have been released which provide valuable information to decipher the molecular biology of these blood flukes. However, progress in identifying and characterizing effective drug targets and vaccine candidates has been severely hampered by a general paucity of suitable molecular tools for modulation of critical genes in schistosomes. RNA interference (RNAi) has been developed as a post-transcriptional gene silencing tool over the past decade for loss-of-function studies in helminths [11–15]. Indeed, many reports have described transient knockdown of gene products using RNAi, but the functional assays used were deployed only *in vitro*, a culture state that generates information on parasites that have been weakened following their removal from the host [11–15].

The clustered regularly interspaced short palindromic repeat (CRISPR) approach has emerged as a novel genomic editing tool that has been broadly adapted in various organisms [16–26]. CRISPR was first identified in bacteria as a defense mechanism against foreign genetic elements, utilizing RNA-guided CRISPR-associated protein (Cas) endonucleases to recognize and cleave invading viral DNA [27–32]. The CRISPR system has been repurposed for transcription regulation using a catalytically dead version of Cas9 (dCas9) (with mutations at H840A and D10A) lacking endonucleolytic activity [33]. The dCas9-sgRNA (single guide RNA) complex can specifically interfere with transcriptional initiation or transcriptional elongation, in a process which was termed CRISPR interference (CRISPRi) [33, 34]. The dCas9 protein can also be fused to transcriptional repressor domains [eg. Krüppel-associated box (KRAB)] to achieve more effective transcriptional silencing [34–37]. CRISPRi is a simple and cost-effective gene regulation tool with greater versatility, higher efficacy and specificity [38–40]. To date, CRISPRi has been broadly employed for transcription repression, directed revolution, metabolic engineering and targeted genetic screening in mammalian cells, in the zebra fish, in *Caenorhabditis elegans* and a variety of unicellular organisms such as *Synechococcus elongatus*, *Saccharomyces cerevisiae*, *Toxoplasma gondii* and *Plasmodium falciparum* [33–45]. To advance more efficient gene regulation in parasitic helminths, we applied CRISPRi in *S. mansoni*.

The life cycle and morphology of schistosomes are both complex. Adult *S. mansoni* pass eggs that escape the mammalian host in faeces. The egg gives rise to a ciliated larva, the *miracidium* that infects its specific *Biomphalaria* snail host, within which it undergoes a dramatic body conversion to produce an obligate asexually-reproducing adult, the mother sporocyst. Endogenous proliferation of stem cells (=germinal cells in the asexual stage) in the mother sporocyst leads to a new asexual stage - the daughter sporocyst [46]. These ‘daughters’, in turn, can generate, by germinal cell proliferation, the next stage, the migratory cercariae, that escape from the snail into the aquatic environment [47]. Cercariae seek and then penetrate the skin of a mammal, transform into schistosomula, which enters the vasculature to develop into dimorphic sexual adults. Intra-mammalian development, driven by stem cells (defined as neoblast cells), gives rise to the remarkable reproductive capability of adults [48], over prolonged periods - indeed, schistosomes can live for over 30 years [49]. To unravel the critical roles of stem cells in driving the schistosome life cycle [50–54], characterization of the somatic stem cell marker SmFGFRA has drawn increased attention [50, 52–56]. Our previous study demonstrated SmFGFRA is abundantly expressed in different *S. mansoni* developmental stages [57]. The distribution pattern of SmFGFRA in embryonic cells of immature eggs, in the neural mass of mature eggs and miracidia, and its co-location with EdU^+^ (thymidine analog 5-ethynyl-2’-deoxyuridine) cells in adult *S. mansoni*, strongly implicated its important roles in maintaining schistosome stem cells, in development of the nervous and reproductive systems, and in the host-parasite interplay [57].

We report, for the first time, the application of CRISPRi in *S. mansoni* targeting the *SmfgfrA* gene. We first pre-screened eight sgRNAs, designed specifically targeting different loci of *SmfgfrA*, for effective inhibition of its transcription initiation and elongation. We determined the distinct phenotypic changes in *SmfgfrA*-repressed adult worms, schistosomula and eggs. Then, we injected the *SmfgfrA*-repressed eggs into the tail vein of mice and explored the pathogenicity induced by these *SmfgfrA*-repressed eggs *in vivo*.

## 2 Methods

### 2.1 Ethics

All experiments were approved by the Animal Ethics Committee (ethics number P242) of the QIMR Berghofer Medical Research Institute. The study was carried out based on the guidelines of the National Health and Medical Research Council of Australia, as published in the Australian Code of Practice for the Care and Use of Animals for Scientific Purposes, 7th edition, 2004 (www.nhmrc.gov.au). All work involving live *S. mansoni* parasites was conducted in quarantine-accredited premises.

### 2.2 Maintenance of Parasites

Swiss mice (female, 6 weeks old) were infected with 100 *S. mansoni* cercariae subcutaneously. The infected mice were euthanized seven weeks post-infection and adult worms were harvested by portal perfusion using 37°C pre-warmed RPMI Medium 1640 (Gibco, Sydney, Australia). Adult worms were cultured overnight in RPMI complete medium [RPMI Medium 1640 (Gibco) supplemented with 10% (v/v) heat-inactivated fetal bovine serum (FBS, Gibco) and 100 IU/ml penicillin and 100 μg/ml streptomycin (Gibco)] at 37°C in an atmosphere of 5% CO_2_. Mouse livers were removed at necropsy and liver eggs were isolated and purified as described [58]. Liver eggs were cultured in RPMI complete medium at 37°C in 5% CO_2_. *S. mansoni* cercariae were obtained by shedding infected *Biomphalaria glabrata* snails under bright light. Schistosomula were obtained by mechanical transformation of cercariae *in vitro* and cultured in Basch’s medium as described [59].

### 2.3 Design of Guide RNA Targets and Reconstruction of Vectors

Single-guide RNAs (sgRNAs) were designed utilizing the web-based tools available at https://bioinfogp.cnb.csic.es/tools/breakingcas/ [60] and the Benchling application (https://benchling.com) to predict binding sites for the *Streptococcus pyogenes* dCas9 nuclease within the genome of *S. mansoni*. The location of the transcriptional start site (TSS) was determined using WormBase ParaSite (https://parasite.wormbase.org/index.html). *SmfgfrA* (Smp_175590) comprises thirteen exons separated by twelve introns spanning 51.95 kb on the reverse strand of *S. mansoni* chromosome 1, including a 64 bp 5’ untranslated region (5’ UTR) (**Figure 1A**). Eight sgRNAs (i1-i8) were designed uniquely targeting either DNA strand of *SmfgfrA* to determine whether CRISPRi could induce efficient repression of transcription initiation/elongation of this gene in *S. mansoni*. SgRNA i1, i2, i3, i7, and i6 align to nucleotides 667-686 (+67 bp to +86 bp relative to the predicted TSS), 703-722 (+103 bp to +122 bp), 707-726 (+107 bp to +126 bp), 744-763 (+144 bp to +163 bp), and 783-802 (+183 bp to +202 bp) in exon 1, respectively (Figure 1B); sgRNA i5 and i4 target nucleotides 871-890 (+271 bp to +290 bp) and 910-919 (+310 bp to +329 bp) in exon 2, respectively (**Figure 1B**). For the inhibition of transcription initiation, sgRNA i8 was designed to target residues 580-599 (−21 bp to −2 bp upstream of the TSS) of *SmfgfrA* (**Figure 1B**). All sgRNAs are adjacent to the protospacer adjacent motif (PAM), NGG. SgRNA oligonucleotides and PAM sequences are listed in **Table 1**. A non-targeting sgRNA (5’-GACCAGGATGGGCACCACCC-3’) was used as a negative control (NC). The sgRNAs were synthesized as double-stranded DNA fragments flanked by BstXI and XhoI restriction sites (Integrated DNA Technologies, Singapore) and inserted into the vector, pgRNA-humanized (a gift from Stanley Qi, Addgene plasmid #44248) [33]. Expression of sgRNA is driven by the mouse U6 promoter. The CRISPRi vector PHR-SFFR-dCas9-BFP-KRAB (a gift from Stanley Qi & Jonathan Weissman, Addgene plasmid #46911) [34] contains a silencing-prone spleen focus forming virus (SFFV) promoter expressing dCas9 fused to the KRAB transcription repressor.

**Figure 1.**
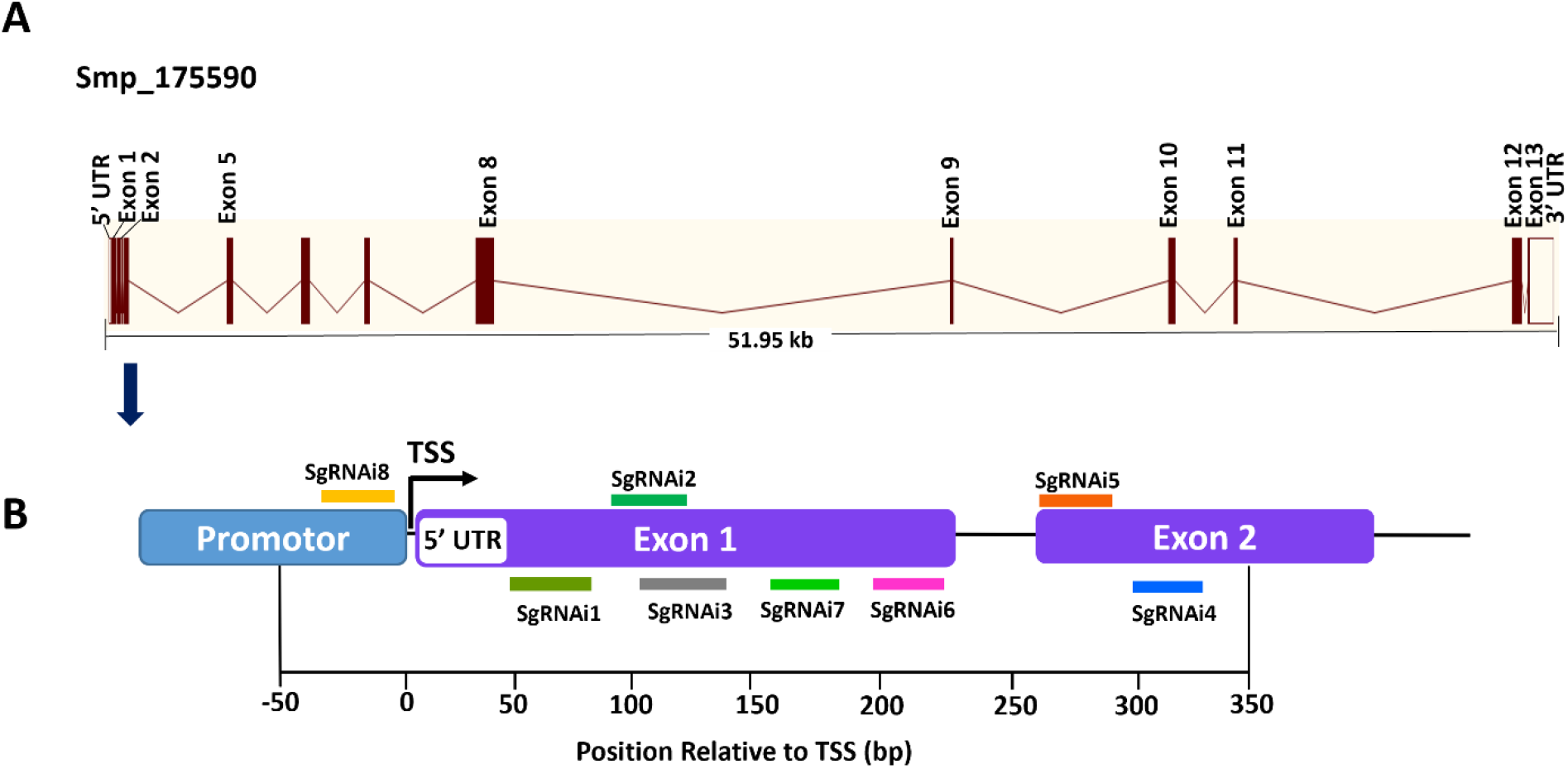
Gene structure of *S. mansoni* fibroblast growth factor receptor A (*SmfgfrA*) and locations of singe-guide RNAs (sgRNAs). (**A**) Schematic diagram of the *SmfgfrA* (Smp_175590) gene containing thirteen exons, twelve introns, 5’ untranslated region (5’ UTR) and 3’ UTR, spanning 51.95 kb on the reverse strand of *S. mansoni* chromosome 1. (**B**) Diagram of *SmfgfrA* indicating target sites of eight sgRNAs (i1-i8). SgRNAs (i1-i7) were designed for inhibition of transcription elongation within a window from +67 bp to +329 bp relative to the transcriptional start site (TSS) of *SmfgfrA*. SgRNA i8 aligns to the promotor region (−21 bp to −2 bp) of *SmfgfrA* for suppression of transcription initiation.

**Table 1.**
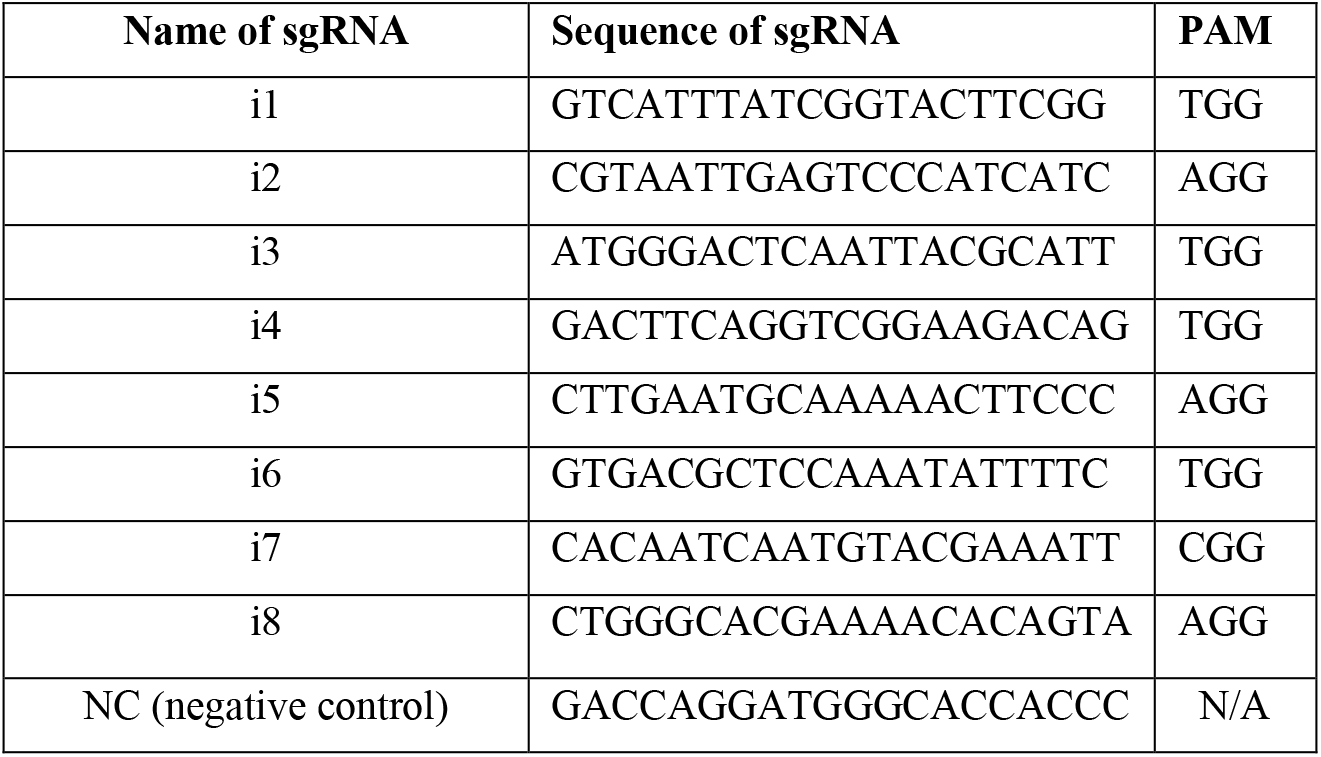
Sequences of sgRNAs.

### 2.4 Transfection of *S. mansoni* Parasites

To perform CRISPRi repression targeting *SmfgfrA*, pools of 5 pairs of adult *S. mansoni* or 10,000 liver eggs or 2000 schistosomula were subjected to electroporation in 200 μl Opti-MEM containing 3 μg PHR-SFFR-dCas9-BFP-KRAB vector and 3 μg of pgRNA vector reconstructed with: (1) Negative control sgRNA (non-targeting RNA sequence) (NC); (2) i1 sgRNA (i1); (3) i2 sgRNA (i2); (4) i3 sgRNA (i3); (5) i4 sgRNA (i4); (6) i5 sgRNA (i5); (7) i6 sgRNA (i6); (8) i7 sgRNA (i7); (9) i8 sgRNA (i8); (10) i4 sgRNA and i5 sgRNA (i4+i5, 3 μg of each); (11) i4 sgRNA+i5 sgRNA+i6 sgRNA (i4+i5+i6, 3 μg of each); or (12) i5 sgRNA+i8 sgRNA (i5+i8, 3 μg of each). The mixture was transferred into a pre-chilled 4 mm electroporation cuvette (Bio-Rad, Sydney, Australia) and subjected to square wave electroporation using a single 20 millisecond pulse of 125 Volts (Gene Pulser Xcell Electroporator, Bio-Rad) [61]. Thereafter, electroporated eggs and adult worms were cultured in RPMI complete medium and schistosomula were cultured in Basch’s medium [2] at 37°C in 5% CO_2_. Wild type (WT) parasites (adult worms or eggs or schistosomula) were not subjected to electroporation but were cultured under the same conditions as control. *SmfgfrA*-modulated parasites and control parasites (WT and NC-treated) were harvested after three days in culture. Hatched miracidia were collected from *SmfgfrA*-modulated eggs and control eggs cultured for seven days post-electroporation and the egg hatching efficiency (%) was determined by dividing the number of hatched eggs by the total number of eggs X 100. Eggs laid *in vitro* by adult female worms were collected three days post-electroporation and the collected eggs in each group were counted.

### 2.5 Real-time PCR

Total RNAs were extracted from *SmfgfrA*-repressed and control (WT and NC-treated) *S. mansoni* eggs, schistosomula, adult worms and miracidia hatched from *SmfgfrA*-repressed and control eggs using RNeasy Mini Kits (Qiagen, Melbourne, Australia), followed by cDNA synthesis using QuantiTect Reverse Transcription Kits (Qiagen). Real-time PCR was conducted using QuantiNova SYBR^®^ Green PCR Kits (Qiagen) on a Mic qPCR Cycler (Bio Molecular Systems, Upper Coomera, QLD, Australia). Forward primer (5’-ATGGGACTCAATTACGCATT-3’) and reverse primer (5’-CACCACTGTCTTCCGACCTG-3’) for *SmfgfrA* were designed using the Primer 3 software (http://frodo.wi.mit.edu/), and the specificity of the primer sequences was confirmed by the Basic Local Alignment Search Tool (BLAST) at the NCBI website (http://blast.ncbi.nlm.nih.gov/Blast.cgi). *S. mansoni* glyceraldehyde-3-phosphate dehydrogenase (GAPDH) was used as the house keeping reference gene [62]. Real-time PCR reactions contained 10 μl 2xSYBR Green PCR Master Mix, 100 ng cDNA, and 0.7 μM of each primer. The cycling parameters were set as follows: 95°C for 5 min, 40 cycles of 95°C for 30 s, 58°C for 30 s and 72°C for 30 s. Data analysis was performed using the Mic qPCR software (Bio Molecular Systems). Relative *SmfgfrA* transcription levels in each group were determined using the 2^-ΔΔCt^ calculation [63] by normalizing to the control parasites (NC-treated parasites).

### 2.6 Caspase-3/-7 Activity Assay

*S. mansoni* soluble egg antigen (SEA), soluble worm antigen preparation (SWAP) and soluble schistosomula native antigens were prepared in PBST (PBS+0.1% v/v Tween-20) plus 10 mM HEPES and cOmplete^™^, Mini, EDTA-free Protease Inhibitor (Sigma-Aldrich, Sydney, Australia) as described [64, 65]. Protein concentration was determined using the Bradford assay as described [66]. The caspase-3/-7 activity of SWAP (0.4 mg/ml), SEA (0.2 mg/ml) and soluble schistosomula antigens (0.2 mg/ml) was determined using the Caspase-Glo^®^ 3/7 Assay System (Promega, Sydney, Australia) according to the manufacturers’ instructions.

### 2.7 Western Blotting

*S. mansoni* SWAP (40 μg) and SEA (20 μg) were separated on 12% (w/v) SDS-PAGE gels and transferred to an Immun-Blot low fluorescence-PVDF membrane (Bio-rad). The membrane was first blocked with Odyssey Blocking Buffer (TBS) (LI-COR Biosciences, Lincoln, Nebraska, USA) for 1 h at room temperature with shaking. Subsequently, the membrane was incubated with a polyclonal mouse anti-rSmFGFRA-L (anti-recombinant SmFGFRA extracellular ligand binding domain) antibody (previously generated in our laboratory [57]) (1:100 diluted in Odyssey buffer with 0.1% (v/v) Tween-20) for 1 h with shaking at room temperature. Following washes (4X) in Tris-Buffered Saline (TBS)/0.1% Tween-20 (TBST), the membrane was incubated with IRDye-labeled 680LT goat anti-mouse IgG antibody (Li-COR Biosciences) (1:15,000 diluted in Odyssey buffer with 0.1% Tween-20 and 0.01% SDS) for 1 h with shaking in a dark chamber. After four washes with TBST, the membrane was dried in the dark and visualized using the Odyssey CLx Infrared Imaging System [65]. An anti-actin antibody (Sigma-Aldrich) was used to probe the expression of actin to ensure the equal loading of samples.

### 2.8 EdU Staining

*SmfgfrA-modulated* parasites were cultured at 37°C under 5% CO_2_ in RPMI complete medium containing 10 μM EdU (Thermo Fisher Scientific, Brisbane, Australia), which specifically stains stem cells in *S. mansoni* [54]. After 24 h, the stained worms were fixed in 10% formalin, paraffin embedded and sectioned. Three sections (4 μm/section, with 4 μm distance between each section) of EdU-labeled adult worms and schistosomula were subjected to EdU detection using a Click-iT^™^ EdU Cell Proliferation Kit (Alexa Fluor^™^ 488 dye) (Thermo Fisher Scientific) according to the manufacturers’ instructions. Nuclei in all tissue sections were also stained with Propidium iodide (PI) (Sigma-Aldrich) and visualized using a Zeiss 780 NLO confocal microscope (Zeiss, Oberkochen, Germany). Counting of EdU^+^ cell nuclei and PI^+^ cells was performed using QuPath software (https://qupath.github.io) [67]. The percentage of EdU-positive cells was calculated by dividing the EdU^+^ cell number with the PI^+^ cell number X 100.

### 2.9 Miracidial Behavioral Assay

Miracidia were hatched in deionized water under light from *SmfgfrA*-repressed and control (WT, NC-treated) eggs. Miracidia were then harvested [59] and their behavior were monitored as described [68, 69]. Briefly, approximately 30 *S. mansoni* miracidia in 100 μl deionized water were distributed evenly to a microscope slide. Miracidial movement (swimming) in the field of view (FOV) was detected utilizing an Olympus-CKX41 microscope equipped with an Olympus DPI Digital Microscope Camera DP22 (25 frames per second at 2.8-megapixel image quality). Miracidial movement was recorded for 1 minute by video and followed by analyzing with the FIJI software to calculate velocity (speed) of miracidial swimming, the tortuosity (the ratio of track length to maximum displacement) of miracidial swimming and the duration (time) of miracidia staying in the FOV [68]. The miracidial movement velocity was determined in pixels employing the rolling mean subtraction approach [68, 69]. The location of miracidial was monitored in each frame along an x-y axis and the trajectories were interpolated utilizing TrackMate (the plugin for FIJI software) [69, 70]. Applying the MTrackJ (ImageJ plugin), the average velocity, duration and tortuosity of miracidia movement in each video were determined. The heatmaps were produced as mentioned [69, 71] for demonstrating the movement pattern of individual miracidia.

### 2.10 Intravenous Injection of *SmfgfrA-repressed* Eggs into Mice

Swiss mice were injected intravenously (i.v.) with *SmfgfrA-repressed* or control (WT eggs and NC-treated eggs) eggs as described [61]. Briefly, eggs were cultured for 48 h after electroporation, followed by three washes with chilled PBS. Then, 1000 eggs in 100 μl sterile PBS were injected into the lateral tail vein of female Swiss mice (8-9 weeks of age). Mice injected with sterile PBS and control eggs (WT eggs and NC-treated eggs) were served as negative control mice. Mice (5 mice/group) were euthanized two weeks post-injection.

### 2.11 Serum IgE level and Lung Granuloma Size in Mice Exposed to *SmfgfrA-repressed* Eggs

Two weeks post-injection, blood was obtained from each mouse and sera were prepared individually. The total immunoglobulin E (IgE) level in the serum samples was measured using an IgE mouse ELISA kit (Thermo Fisher Scientific, Waltham, Mass, USA) according to the manufacturer’s instructions.

To determine the size of granuloma that had formed in the lungs, the left lung of each mouse was fixed in 10% (v/v) formalin, paraffin embedded and sectioned. Sections (4 μm) of these paraffin blocks were stained with Hematoxylin and Eosin (H&E) to evaluate inflammatory infiltrates and the cellularity of granulomas. Slides were imaged using an Aperio Slide Scanner (Aperio Technologies, Vista, CA, USA) and analyzed using Aperio Image Scope v11.1.2.760 software (Leica Biosystems Imaging, Buffalo Grove, IL, USA). The degree of lung pathology was quantified by measurement of the area density of granulomatous lesions. The granuloma ratio in each lung sample was estimated from the total area of the granulomas in the lung sample divided by the total area of the lung tissue and X 100. For each group, 10 slides per lung (5 lungs/group), totaling 50 samples, were measured.

### 2.12 Statistical Analysis

GraphPad Prism software (Version 8.2.1, La Jolla, CA, USA) was used for all statistical analyses. All data are shown as the mean ± SE. Differences between groups were analyzed for statistical significance by One-way ANOVA and, where appropriate, by two-tailed Student’s t-test. A statistically significant difference for a particular comparison was defined as a *p* value ≤ 0.05. \**p* value≤ 0.05, \*\**p* value≤ 0.01, \*\*\**p* value ≤ 0.001, **** *p* value ≤ 0.0001, not significant (ns).

## 3 Results

### 3.1 CRISPRi-mediated Repression of *SmfgfrA* in *S. mansoni* Adult Worms

#### *SmfgfrA* Transcription Level in *SmfgfrA-repressed* Adult Worms

The transcription level of *SmfgfrA* was quantified in CRISPRi-modulated *S. mansoni* adult worms, liver eggs and schistosomula using real-time PCR assays. We found the most reduced level of transcription of *SmfgfrA* was induced by sgRNA i5 (51.6%, *p*<0.0001) in adult worms, followed by i4 (43.7%, *p*<0.0001), i6 (27.7%, *p*=0.0018), i7 (22.6%, *p*=0.0083), and i8 (15.4%, *p*=0.0184) (**Figure 2A)**. No significant downregulation of *SmfgfrA* transcripts was observed in worms electroporated with i1, i2 or i3, compared with the control groups (WT and treated with NC) (**Figure 2A**). As a result, we selected i4 and i5 for subsequent study. To further investigate whether the gene repression efficiency was sex dependent, we determined *SmfgfrA* transcription levels in i4-treated and i5-treated male and female worms. Notably, both sgRNA i4 and sgRNA i5 induced similar transcriptional reductions in *SmfgfrA* in both male and female worms (**Figure 2B**); accordingly, we used paired adult worms in all subsequent studies.

**Figure 2.**
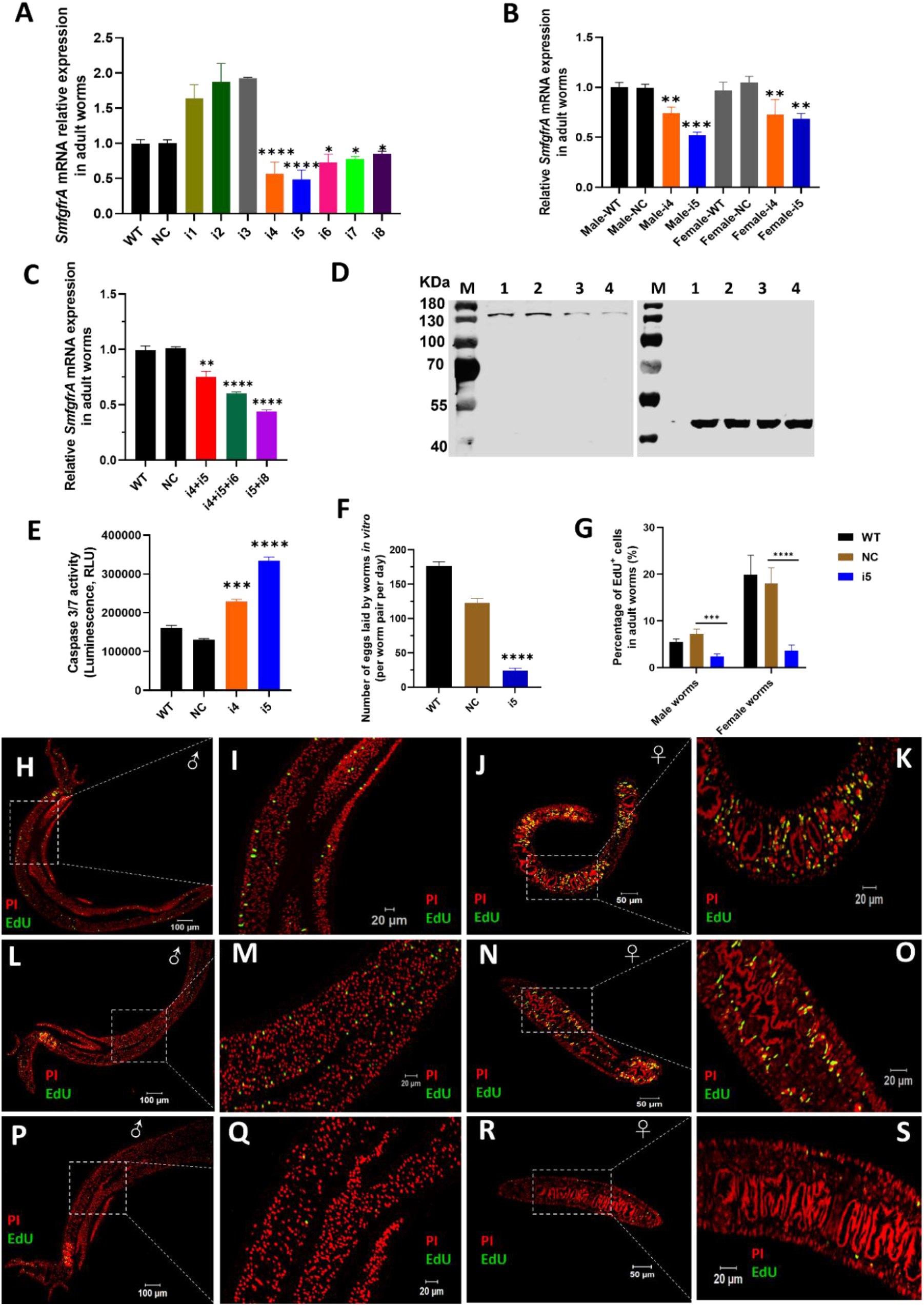
CRISPRi of *SmfgfrA* in *S. mansoni* adult worms. Transcription level of *SmfgfrA* was determined in (**A**) paired adult worms electroporated with a CRISPRi vector (PHR-SFFR-dCas9-BFP-KRAB) combined with reconstructed pg-RNA vector containing sgRNA (i1, i2, i3, i4, i5, i6, i7, i8), individually; (**B**) adult male and female worms treated with sgRNA i4 or i5; (**C**) paired worms treated with sgRNA i4+i5, i4+i5+i6, or i5+i8. Wild-type (WT) worms and worms treated with CRISPRi vector combined with pg-RNA vector containing a negative control, scrambled sgRNA sequence (NC) were used as controls. (**D**) Western blot demonstrating expression level of SmFGFRA protein in soluble worm antigen preparation (SWAP) of WT adult worms (Lane 1) and worms treated with NC (Lane 2), i4 (Lane 3) or i5 (Lane 4). An anti-actin antibody was employed to ensure equal protein loading. The electrophoresed SWAPs were transferred for the western blot analysis to two PVDF membranes which were probed simultaneously with the anti-rSmFGFRA-L antibody (left blot) and the anti-actin antibody (right blot). (**E**) Caspase-3/-7 activity measured in i4 and i5-treated adult worms. This assay was conducted in duplicate with all data presented as the mean ± SE. (**F**) The number of eggs laid by WT adult worms, NC-treated adult worms or i5-treated adult worms *in vitro*. (**G**) The effect of *SmfgfrA* repression on the percentage of EdU^+^ cells in i5-treated male worms and female worms. WT and NC-treated male and female worms were used as controls. Cell nuclei in all samples were also stained with Propidium iodide (PI). Percentage of EdU^+^ cells in each sample was calculated by dividing the EdU^+^ cell number with PI^+^ cell number X 100. All data are shown as the mean ± SE (WT male worm: n=13, WT female worm: n=8, NC male worm: n=12, NC female worm: n=8, i5 male worm: n=10, i5 female worm: n=15). Confocal projections representative signals of EdU (green) and PI (red) in (**H**) WT male worm, (**J**) WT female worm, (**L**) NC-treated male worm, (**N**) NC-treated female worm, (**P**) i5-treated male worm and (**R**) i5-treated female worm. (**I**) (**K**) (**M**) (**O**) (**Q**) (**S**) are magnified squared-region in (H) (J) (L) (N) (P) (R), respectively. (Statistical significance was established employing One-way ANOVA by comparing with NC group: * *p* value≤ 0.05, \*\*\**p* value ≤ 0.001).

It has been shown that CRISPRi conducted in *Escherichia coli* (*E. coli*) using multiplexed sgRNAs targeting the same gene can markedly increase gene repression efficiency compared with using single sgRNAs [33]. To investigate this scenario in schistosomes, we tested the combination of sgRNA i4+i5, i4+i5+i6 and i5+i8, and this resulted in the downregulation of gene transcription by 25.6% (*p*=0.0017), 40.25% (*p*<0.0001) and 56.5% (*p*<0.0001) in the treated adult *S. mansoni* worms respectively (**Figure 2C**), compared with NC-treated parasites. Given the similar gene repression efficiency induced by the CRISPRi using single sgRNAs or multiplexed sgRNAs in silencing *SmfgfrA*, we decided to perform CRISPRi with individual sgRNA i5 and/or sgRNA i4 for subsequent phenotypic change studies.

#### Reduction of SmFGFRA Protein Expression in *SmfgfrA-repressed* Adult Worms

To determine whether the repression of *SmfgfrA* was reflected at the translational level, we performed western blot analysis using an anti-rSmFGFRA-L polyclonal antibody to probe SWAP extracted from CRISPRi-modulated adult worms. SEA generated from WT worms and NC-treated worms served as controls. A clearly decreased level of SmFGFRA protein expression was evident in SWAP extracted from i4-treated (**Figure 2D**, Lane 3) and i5-treated (**Figure 2D**, Lane 4) adult worms compared with that from WT adults (**Figure 2D**, Lane 1) and NC-treated adults (**Figure 2D**, Lane 2).

#### Increased Caspase-3/-7 Activity in *SmfgfrA-repressed* Adult Worms

Caspase-3 and caspase-7 are major executioner caspases that play critical roles in coordinating cell apoptosis, and thus determining their activities has been broadly used for monitoring apoptosis [72–74]. Therefore, we determined the effect of *SmfgfrA*-repression on apoptosis in extracts of CRISPRi-modulated parasites by measuring the activity of caspase-3/-7. Remarkably, caspase-3/-7 activity in i4-treated and i5-treated adult worms was enhanced by 43% (*p*=0.0003) and 60.9% (*p*<0.0001), respectively (**Figure 2E**).

#### Reduced Adult Worm Egg Production

To further investigate the effects of *SmfgfrA*-repression on egg production by adult females, the number of eggs laid *in vitro* by CRISPRi-modulated paired worms was determined. Notably, the egg production in i5-treated adult worms was dramatically depleted by 80.4% (*p*<0.0001) compared with NC-treated worms (**Figure 2F**).

#### Decline in EdU Incorporation in *SmfgfrA-repressed* Adult Worms

EdU is able to be incorporated into newly synthesized cellular DNA, and stains only proliferating stem cells in schistosomes [54]. To assess the effect of *SmfgfrA-repression* on schistosome stem cells, we monitored EdU incorporation in CRISPRi-modulated parasites. Representative confocal images of EdU signaling in sections of adult males and females are shown in **Figure 2H–2S**. We found that ~7.17% and ~18% of cells in NC-treated male and NC-treated female worms, respectively, were EdU^+^, whereas the percentage of EdU^+^ cell nuclei was markedly reduced to ~2.53% (decreased by 64.7%, *p*=0.0007) in i5-treated male worms, and to ~3.8% (reduced by 78.9%, *p*<0.0001) in i5-treated female worms (**Figure 2G**).

### 3.2 CRISPRi-induced Silencing of *SmfgfrA* in *S. mansoni* Schistosomula

In CRISPRi-modulated schistosomula, we found *SmfgfrA*-specific transcripts were clearly repressed by 17.9% (*p*=0.0006) in i5-treated parasites, while no significant change in the *SmfgfrA* mRNA level was observed in i4-treated schistosomula (**Figure 3A**). We then examined apoptosis in schistosomula following repression of *SmfgfrA* by measuring caspase-3/-7 activity. In i5-transfected schistosomula, the caspase-3/-7 activity was clearly increased by 21.7% (*p*=0.0057) but no significant difference was evident in i4-treated schistosomula, compared with the NC group (**Figure 3B**). EdU labelling was undertaken to determine whether the silencing of *SmfgfrA* affected germinal cells in schistosomula. **Figure 3D-F** shows confocal images of EdU signal in sections of *SmfgfrA-repressed* schistosomula. We found the percentage of EdU^+^ cells in i5-treated schistosomula (~2.3% EdU^+^ cells) was considerably reduced by 45.6% (*p*=0.005) when compared with NC-treated schistosomula (~4.3% EdU^+^ cells) (**Figure 3C**).

**Figure 3.**
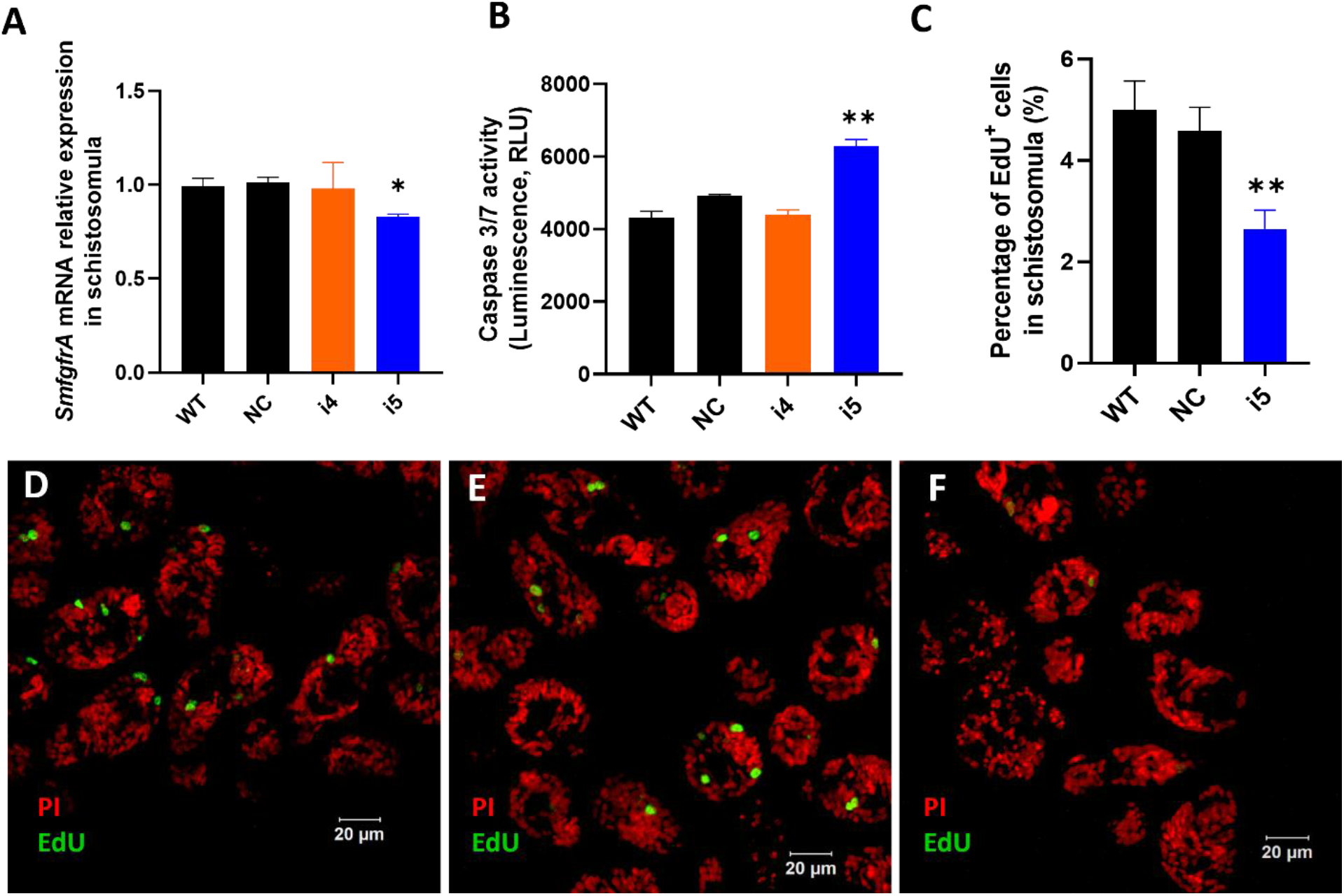
CRISPRi-mediated *SmfgfrA* repression in *S. mansoni* schistosomula. (**A**) *SmfgfrA* transcription level in untreated WT schistosomula and schistosomula treated with NC, i4 or i5. (**B**) Effect of *SmfgfrA-repression* on caspase-3/-7 activity in i4-treated or i5-treated schistosomula. WT schistosomula and schistosomula treated with NC were used as controls. Experiments in panels (A)-(B) was performed in duplicate with data presented as the mean ± SE. (**C**) The percentage of EdU^+^ cell nuclei in WT schistosomula and schistosomula treated with NC, i4 or i5. All data are demonstrated as the mean ± SE (WT: n=33; NC: n=42; i5: n=42). Confocal projections representing EdU (green) and PI (red) labeled (**D**) WT schistosomula, (**E**) NC-treated schistosomula and (**F**) i5-treated schistosomula.

### 3.3 CRISPRi-induced Repression of *SmfgfrA* in *S. mansoni* Eggs

#### Reduced *SmfgfrA*-specific Transcripts

*SmfgfrA* transcription levels in CRISPRi-repressed eggs treated with i4 and i5 were substantially reduced by 46.9% (*p*=0.0329) and 67.3% (*p*=0.0056), respectively (**Figure 4A**). Notably, a marked decrease (25.3%, *p*=0.0025) of *SmfgfrA*-specific transcripts was also detected in miracidia hatched from i5-treated eggs whereas no significant change was evident in miracidia collected from i4-treated eggs (**Figure 4B**).

**Figure 4.**
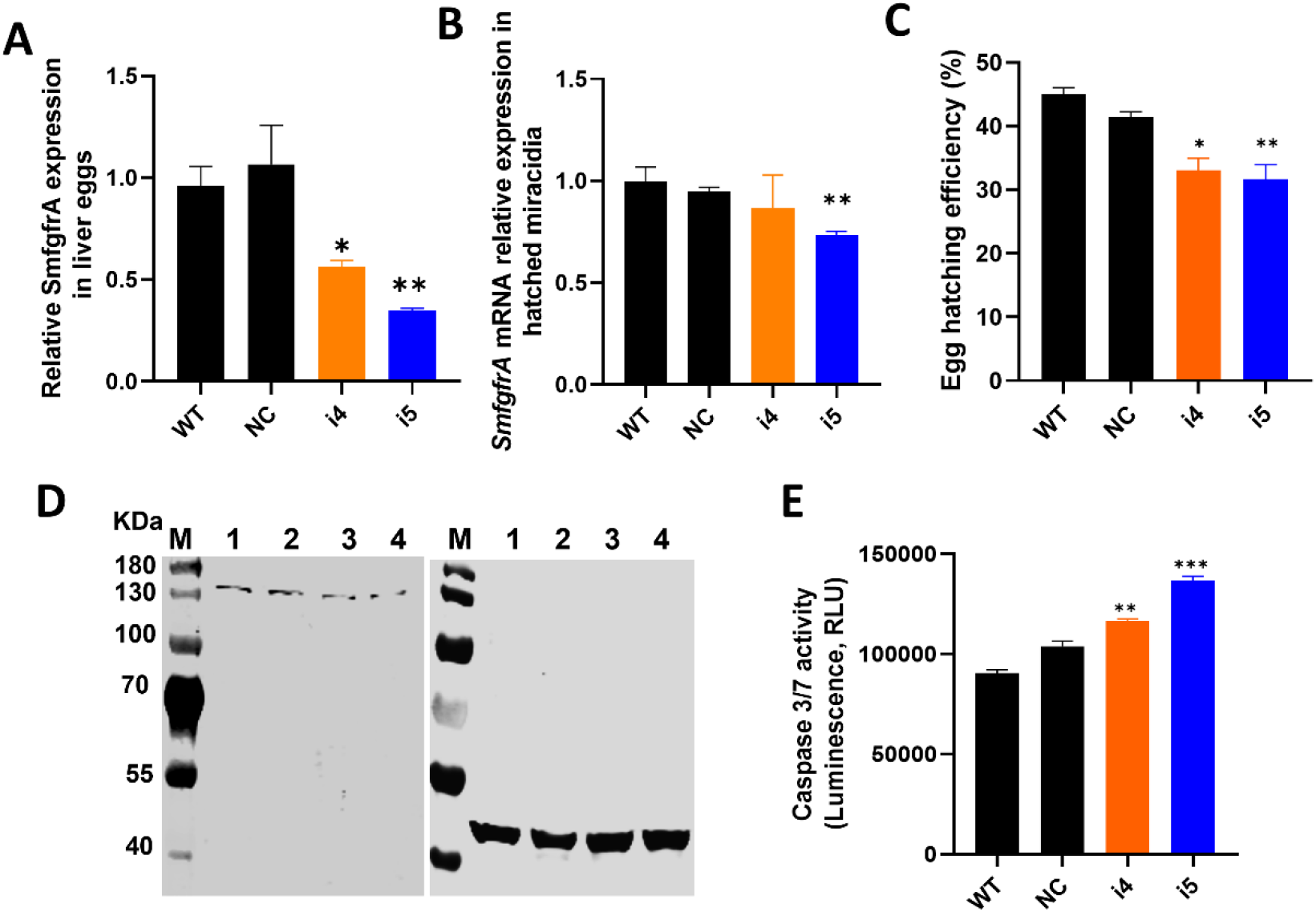
CRISPRi of *SmfgfrA* in *S. mansoni* eggs. (**A**) *SmfgfrA* mRNA level in eggs treated with i4 or i5. WT eggs and NC treated eggs were used as controls. (**B**) Transcription level of *SmfgfrA* in miracidia hatched from WT eggs and eggs treated with NC, i4 or i5. (**C**) Effects of *SmfgfrA* repression on the hatching of *S. mansoni* eggs. The hatching efficiency (%) of WT eggs and eggs treated with NC, i4 or i5 was calculated by dividing the number of hatched eggs with the total number of eggs (hatched and unhatched) X 100. Experiments in panels (A-C) was performed in triplicates with all data presented as the mean ± SE. (**D**) Caspase-3/-7 activity in soluble egg antigen (SEA) of WT eggs and eggs treated with NC, i4 or i5. This assay was conducted in duplicate with all data presented as the mean ± SE. (**E**) Western blot showing expression level of SmFGFRA protein in SEA of WT eggs (Lane 1) and eggs treated with NC (Lane 2), i4 (Lane 3) or i5 (Lane 4). An anti-actin antibody was utilized as control to ensure equal loading. The electrophoresed SEAs were transferred for the western blot analysis to two PVDF membranes which were probed simultaneously with the anti-rSmFGFRA-L antibody (left blot) and the anti-actin antibody (right blot).

#### Decreased Hatching Efficiency of *SmfgfrA-repressed* Eggs

To determine whether *SmfgfrA* suppression had an effect on the hatching ability of modulated eggs, we assessed the hatching ability of WT eggs and eggs treated with NC, i4 or i5. Markedly reduced egg hatching efficiency was evident in i4-treated eggs (20.2%, *p*=0.0156) and i5-treated eggs (23.5%, *p*=0.0069) (**Figure 4C**), compared with eggs treated with NC.

#### Decreased SmFGFRA Expression and Increased Apoptosis in Modulated Eggs

To determine the extent of gene regulation at the protein level, we extracted SEA from *SmfgfrA-* repressed eggs and performed western blotting and caspase-3/-7 activity assays. Consistent with the observations with adult worms, SmFGFRA expression in i4-treated (**Figure 4D**, Lane 3) and i5-treated (**Figure 4D**, Lane 4) eggs was clearly reduced, in comparison with that of NC-treated eggs (**Figure 4D**, Lane 2). Furthermore, caspase-3/-7 activity in i4-treated and i5-treated eggs was elevated by 11% (*p*=0.0097) and 24% (*p*=0.0002) compared to NC-treated eggs, respectively (**Figure 4E**).

#### Modified Behavior of Miracidia Hatched From *SmfgfrA-repressed* Eggs

Behavioral changes of miracidia hatched from *SmfgfrA*-repressed eggs were investigated by analyzing recordings of miracidial movement tracks. Heatmaps were created to illustrate the behavior of individual miracidia within a 1 minute recording. The heatmaps of control miracidia (WT miracidia and miracidia hatched from NC-treated eggs) depicted linear soft blue lines, indicating these miracidia had less circular and faster movement (**Figure 5A - B**). In contrast, there were more circular lines and more abundant red and yellow regions in heatmaps of miracidia hatched from the i5-treated eggs (**Figure 5C**), suggesting more turning and circling behavior and relatively slower swimming of these miracidia. Data analysis showed the swimming velocity of miracidia hatched from i5-treated eggs was markedly decreased by 11.7% (*p*=0.0001) (**Figure 5D**). Notably, the average duration time and the movement tortuosity of miracidia hatched from i5-treated eggs were enhanced by 18.5% (*p*=0.0031) (**Figure 5E**) and 18.9% (*p*=0.003) (**Figure 5F**), respectively.

**Figure 5.**
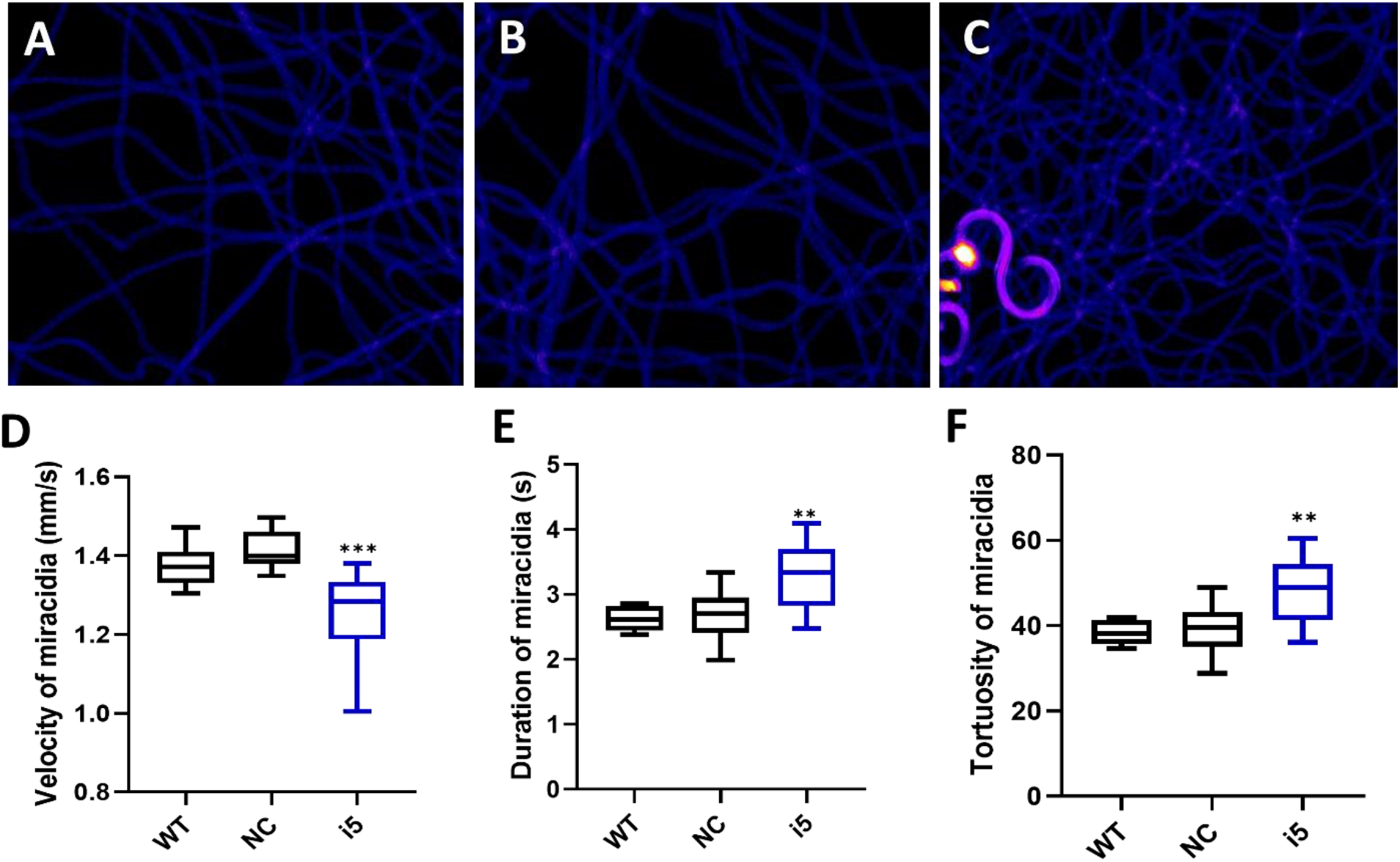
Behavioral changes in *S. mansoni* miracidia hatched from *SmfgfrA*-suppressed eggs. The behavior of miracidia hatched from WT eggs and eggs treated with NC or i5 was monitored. Heatmaps (**A**)-(**C**) represent the movement patterns of individual miracidia within a 1 min recording. Colors in the heatmaps show the time miracidia spent at a specific position. Black: absence; Blue: shorter time presence; Yellow and Red: longer time presence. Boxplots showing (**D**) velocity (**E**) duration, and (**F**) tortuosity of miracidial movement. Experiments were performed in biological duplicates and five technical repeats (n=10).

#### Depleted Granulomatous Inflammation in Lungs of Mice Injected with *SmfgfrA-repressed* Eggs

To determine whether the repression of *SmfgfrA* in eggs could affect the formation of egg-induced granulomas *in vivo*, i5-treated eggs and controls (PBS, WT eggs and NC-treated eggs) were i.v. injected into the lateral vein of the tail of Swiss mice. In mice, schistosome eggs are transported to the lungs via the circulation leading to the development of lung granuloma [61, 75]. Mice were euthanized two weeks post-injection and the left lung of each mouse was harvested for histological analysis. Representative digital microscopic images of mouse lung sections are shown in **Figure 6A-D** and indicate that more intense and severe granulomatous inflammation was evident in lungs of mice injected with WT eggs (**Figure 6B**) and NC-treated eggs (**Figure 6C**) compared with those of mice injected with i5-treated eggs (**Figure 6D**). Calculations of the granuloma ratio from individual lung tissue clearly showed that the sizes of lung granuloma in mice injected with i5-treated eggs were reduced by 60.8% (*p*=0.0003), compared with those in mice injected with NC-treated eggs (**Figure 6E**). Substantially smaller sized granulomas (*p*<0.0001) were also observed in mice injected with NC-treated eggs compared with those present in mice receiving WT eggs (**Figure 6E**), indicating that the electroporation of NC may have subsequently affected the granuloma formation occurred around eggs as previously reported [61].

**Figure 6.**
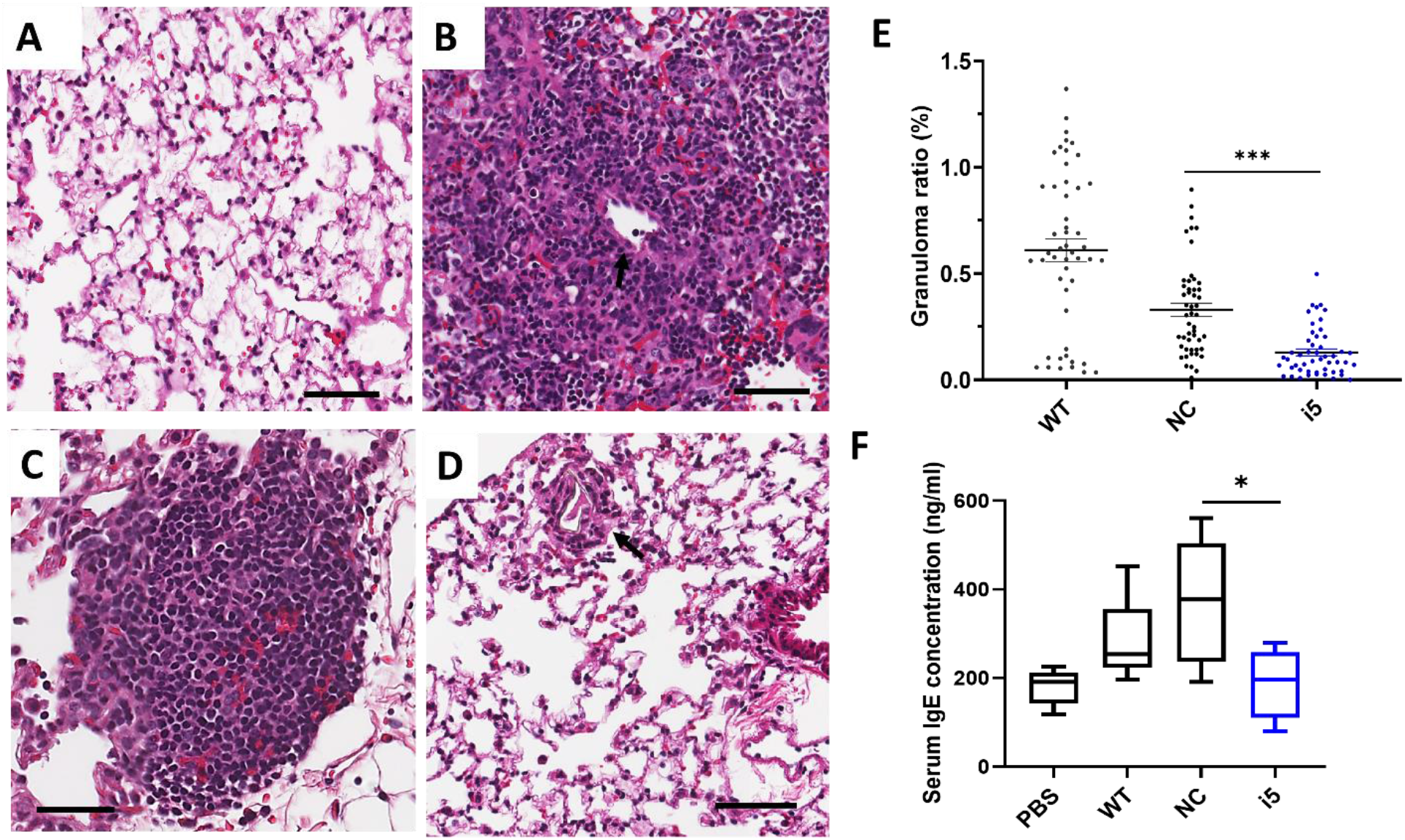
Attenuated granulomatous inflammation and sera IgE levels in mice infected with *SmfgfrA*-repressed schistosome eggs. Representative digital microscopic images showed (**A**) lung from mouse injected with PBS and granulomas in lungs from mice injected with (**B**) WT eggs, (**C**) NC-treated or (**D**) i5-treated eggs. All sections were H&E stained. Scale bars in (A) - (D) = 50 μm. (**E**) Effect of *SmfgfrA*-repressed eggs on granuloma formation in the lungs of mice injected with i5-treated eggs. Mice in control groups were infected with untreated WT eggs or NC-treated eggs. Granuloma ratio in each lung section was calculated by dividing the total granuloma size with the lung size X100. Each group included 5 mice and 10 sections per lung were measured (n=50). All data are shown as the mean ± SE. (**F**) Total serum IgE level in mice 2 weeks after i.v. injection of i5-treated eggs. Mice injected with PBS, WT eggs or NC-treated eggs served as controls. This experiment was conducted in duplicate.

#### Reduced Serum IgE Level in Mice Exposed to *SmfgfrA*-repressed Eggs

It is well recognized that during schistosome infection, the host immune response is highly Th2-polarized following egg laying [76]. As a marker of the Th2-polarized response during infection, IgE is associated with the protective response against schistosomes by mediating macrophage toxicity [77]. To determine whether *SmfgfrA* repression of eggs has any effect on the IgE response in infected mice, the levels of IgE were determined in sera collected from mice injected with PBS, WT eggs, NC-treated eggs and i5-treated eggs. Notably, the concentration of serum IgE in mice injected with i5-treated eggs was markedly reduced by 49.8% (*p*=0.00378) compared with that of mice injected with NC-treated eggs (**Figure 6F**).

## 4 Discussion

Since the award of the Nobel Prize in Chemistry (2020) for the discovery of the revolutionary CRISPR/Cas9 gene editing tool, advances in CRISPR technology have accelerated and the approach provides a powerful new avenue to undertake functional genomics studies. Recently, CRISPR/dCas9 has been repurposed for genome regulation instead of genome editing, providing a platform for RNA-guided transcription regulation called CRISPR interference (CRISPRi). Using *S. mansoni* as a model, we show, for the first time, the feasibility of applying CRISPRi to selectively perturb genes in a parasitic worm, and our study points the way forward to undertake loss-of-function studies in schistosomes and other parasitic helminths. Efficient CRISPRi-mediated repression of *SmfgfrA* (the gene encoding the SmFGFRA stem cell marker) was evidenced by the marked downregulation of *SmfgfrA* transcription and a decrease in the expression of SmFGFRA. These features were accompanied by distinct *in vitro* and *in vivo* phenotypic changes including a reduction in the number of stem cells and an elevated level of cell apoptosis in modulated parasites, and the decreased capacity of modulated eggs to induce granulomatous inflammation in infected mice together with a decline in serum IgE levels.

To adapt CRISPRi for application in *S. mansoni*, we started with the screening of 8 sgRNAs (i1-i8) targeting −21 bp to +329 bp relative to the TSS of *SmfgfrA*, through identifying the efficiency of phenotypic changes induced by CRISPRi-*SmfgfrA* modulation in adult *S. mansoni*. Selection of the sgRNA targeting window for *SmfgfrA* was based on a previous study showing that the most effective sgRNAs for CRISPRi in mammalian cells target a region −50 bp to +300 bp relative to the TSS [78]. Consistent with studies in mammalian cells and *E. coli* [33, 40, 79], we found that inhibition of both elongation and initiation of the *SmfgfrA* target gene via CRISPRi were achievable in schistosomes. Notably, the highest efficacy of repression (56.5%) was obtained when CRISPRi was performed by simultaneous recruiting sgRNA i5 and the sgRNA i8. This may indicate a relatively higher gene regulation efficiency occur when inhibiting the target gene elongation and initiation simultaneously in schistosomes, a feature which is worthy of further exploration. Considering the similar gene regulation efficiency of CRISPRi between using single sgRNA i5 and multiplexed sgRNAs i5+i8 in silencing *SmfgfrA*, we selected sgRNA i5 in subsequent phenotypic change studies.

The most reduced level of *SmfgfrA* transcription was observed in *SmfgfrA-*repressed eggs (67.3%), followed by adult worms (51.6%), and the lowest efficiency occurred in schistosomula (18.1%).

A possible reason is that *SmfgfrA* is more highly expressed in eggs than in adult worms and schistosomula [57], resulting in the chromatin in eggs being more open (euchromatin) and providing greater access to dCas9 binding and functioning [80]. Also, the morphology of eggs is relatively simple with a high stem-like cell content [81], compared with other developmental stages, which may indicate increased potential for manipulating the stem cells and stem cell marker genes (eg. *SmfgfrA*) in schistosomes. Furthermore, we found the CRISPRi-induced gene repression efficiency in adult worms is much higher than that observed in schistosomula, although adult worms and schistosomula present similar level of *SmfgfrA* transcripts [57]. This might be explained by the more abundant distribution of this molecule in the tegument of adult worms than that in schistosomula [57], which may result in a relatively higher efficacy of *SmfgfrA*-repression in adult worms as genes with more tegument location may provide more proximity to the external environment containing CRISPR components. In addition, the relatively larger surface area / volume ratio in adult worms than schistosomula [82] may also lead to a higher efficacy of delivering CRISPR components into adult flukes by electroporation.

It is noteworthy that the downregulated transcription of *SmfgfrA* was also detectable in miracidia hatched from *SmfgfrA*-repressed eggs, although the silencing effect declined in these hatched miracidia compared with that observed in modulated eggs. This might be because the eggs (obtained from the livers of mice infected with *S. mansoni*) used for CRISPRi are a mix of immature and mature eggs. The different morphology of these eggs [81, 83] may lead to different level of gene suppression efficiency [84]. Therefore, it is possible that eggs with relatively higher gene silencing efficiency were not able to hatch, which was also evidenced by the decreased hatching efficiency of *SmfgfrA*-repressed eggs, resulting in the silencing effect being reduced in the hatched miracidia. The repression of *SmfgfrA* in miracidia hatched from *SmfgfrA*-modulated eggs was further confirmed by their significantly modified movement behavior in water, represented by enhanced turning and circling movement. This re-emphasizes the importance of *SmfgfrA* in the neuronal functioning and development of schistosomes, a characteristic explored in our previous study showing that SmFGFRA was abundant in the neural mass of mature eggs and miracidia [57]. Together, these outcomes may indicate a long-term effect of CRISPRi-mediated gene repression in schistosomes, as described for *Toxoplasma gondii* [44].

Given the vital role of *SmfgfrA* in maintaining stem cells [50, 54, 57, 85, 86], we examined EdU incorporation in CRISPRi-*SmfgfrA-*modulated adult worms and schistosomula. Consistent with the decrease in *SmfgfrA* transcription levels in these two parasite stages, markedly reduced numbers of EdU^+^ stem cells were identified in both *SmfgfrA*-repressed adult worms and schistosomula, again highlighting the critical roles of *SmfgfrA* in maintaining schistosome stem cells. We also noted that suppression of *SmfgfrA* led to a greater reduction of EdU signal in female worms compared to male worms despite having similar levels of *SmfgfrA*-repressed transcripts. This implies that silencing of *SmfgfrA* may restrain the pairing of male and female worms, or negatively affect the development of the female reproductive system, leading to the pronounced reduction in the number of stem cells present in repressed females. Notably, a remarkably decreased egg production was observed in *SmfgfrA*-repressed worms, indicating the critical role of *SmfgfrA* in reproduction system of this parasite. This was also evidenced by the abundant expression of *SmfgfrA* in vitelline cells [57] which occupy the majority of the female worm and without them, the eggs will not form [1]. Furthermore, the enhanced caspase-3/-7 activity, indicative of increased cell apoptosis, observed in *SmfgfrA*-repressed adult worms, schistosomula and eggs suggests that silencing of *SmfgfrA* may trigger apoptosis in the stem cells of schistosomes. This may be because schistosomes possess mechanisms to eliminate abnormal stem cells via apoptosis, as has been observed in mammalian stem cells [87, 88].

The potentially life-threatening pathology of schistosomiasis is evoked by schistosome eggs that are lodged in mammalian host tissue which induce granulomatous inflammation around the transiting eggs [89]. During a mature schistosome infection when eggs are produced, the immune response of the host is polarized to a Th2-type immunity. Together with this enhancement of the Th2 response, serum IgE levels (a Th2 response marker [77]) and the number of circulating eosinophils are also considerably elevated [77, 90]. To determine whether CRISPRi-mediated suppression of *SmfgfrA* affected the ability of eggs to polarize the Th2 response *in vivo*, we injected *SmfgfrA*-repressed eggs and control eggs into mice and examined the level of serum IgE. The considerably reduced level of IgE observed in mice injected with *SmfgfrA-repressed* eggs indicated decreased Th2 polarization in these animals compared with mice injected with NC-treated eggs. That was further reflected by the decreased sizes of pulmonary circumoval granulomas around *SmfgfrA*-repressed eggs in the lungs of the mice, a feature characteristic of fewer host immune cells (eosinophils, macrophages and neutrophils) accumulating around the modulated eggs [1].

Collectively, these outcomes emphasize the critical role played by SmFGFRA in the formation of schistosome egg-induced lung granulomatous inflammation. This concept is also supported by the abundant distribution of SmFGFRA in the von Lichtenberg’s layer of *S. mansoni* eggs [57], a layer heavily involved in the release of immunogenic secretions through eggshell pores into the host circulatory system which regulate the host immune response and subsequent granuloma formation [91, 92].

These outcomes indicate the feasibility of applying CRISPRi for effective, targeted transcriptional repression in schistosomes, but challenges remain and further additional refinements and optimization of the approach will need to be considered before this revolutionary technique can be applied on a larger scale in parasitic helminths. Strategies developed for other organisms, including utilizing different dCas9 orthologues such as CRISPR1 dCas9 derived from *Streptococcus thermophilus* (*Sth*. dCas9), may improve gene regulation efficiency, a strategy that has already been undertaken in bacteria [44, 45]. Another tactic could be the employment of an alternative transcription repressor domain, such as KRAB-MeCP2, which has been shown to improve gene suppression outcomes in mammalian cells [93].

In conclusion, these findings provide a blueprint for selectively perturbing gene expression by employing, for the first time, CRISPRi in schistosomes with the potential of the approach to be extended to the study of other parasitic helminths, with appropriate refinements and modifications of the methodology to ensure appropriate levels of gene regulation efficacy. With its adaptability and scalability to target diverse gene loci, CRISPRi can provide the basis to markedly accelerate the functional characterization of not only the genomes of schistosomes, but also a range of other parasitic helminths as well.

## DATA AVAILABILITY STATEMENT

The original contributions presented in the study are included in the article. Further inquiries can be directed to the corresponding author.

## AUTHOR CONTRIBUTIONS

Conceived and designed the experiments: H.Y., X.D., D.P.M., J.D.F. and H.S. Performed the experiments: X.D., N.C., and S.R.M. Analyzed the data: X.D., H.Y., D.P.M., C.E.F., and M.K.J. Contributed reagents/materials/analysis tools: X.D. and H.Y. Wrote the paper: X.D., H.Y., D.P.M., J.D.F. and M.K.J.

## ACKNOWLEDGMENTS

We thank Mary Duke from QIMR Berghofer Medical Research Institute for the maintaining of the *S. mansoni* life cycle and the provision of parasite materials for this study. *B. glabrata* snails were provided by the NIAID Schistosomiasis Resource Center of the Biomedical Research Institute (Rockville, MD) through NIH-NIAID Contract HHSN272201700014I for distribution through BEI Resources.

## References

1. McManus, D.P., et al., Schistosomiasis (Primer). Nature Reviews: Disease Primers, 2018.

2. Gryseels, B., et al., Human schistosomiasis. The Lancet, 2006. 368(9541): p. 1106–1118.

3. McManus, D.P., et al. Schistosomiasis—from immunopathology to vaccines. in Seminars in immunopathology. 2020. Springer.

4. Deol, A.K., et al., Schistosomiasis—assessing progress toward the 2020 and 2025 global goals. New England Journal of Medicine, 2019. 381(26): p. 2519–2528.

5. Berriman, M., et al., The genome of the blood fluke Schistosoma mansoni. Nature, 2009. 460(7253): p. 352.

6. Protasio, A.V., et al., A systematically improved high quality genome and transcriptome of the human blood fluke Schistosoma mansoni. PLoS neglected tropical diseases, 2012. 6(1).

7. Zhou, Y., et al., The Schistosoma japonicum genome reveals features of host-parasite interplay. Nature, 2009. 460(7253): p. 345.

8. Luo, F., et al., An improved genome assembly of the fluke Schistosoma japonicum. PLoS neglected tropical diseases, 2019. 13(8).

9. Young, N.D., et al., Whole-genome sequence of Schistosoma haematobium. Nature genetics, 2012. 44(2): p. 221.

10. Stroehlein, A.J., et al., High-quality Schistosoma haematobium genome achieved by single-molecule and long-range sequencing. GigaScience, 2019. 8(9): p. giz108.

11. Dalzell, J.J., et al., Considering RNAi experimental design in parasitic helminths. Parasitology, 2012. 139(5): p. 589–604.

12. Correnti, J.M., P.J. Brindley, and E.J. Pearce, Long-term suppression of cathepsin B levels by RNA interference retards schistosome growth. Molecular and biochemical parasitology, 2005. 143(2): p. 209–215.

13. Fanelli, E., et al., Analysis of chitin synthase function in a plant parasitic nematode, Meloidogyne artiellia, using RNAi. Gene, 2005. 349: p. 87–95.

14. Rinaldi, G., et al., Development of functional genomic tools in trematodes: RNA interference and luciferase reporter gene activity in Fasciola hepatica. PLoS neglected tropical diseases, 2008. 2(7).

15. Vastenhouw, N.L., et al., Long-term gene silencing by RNAi. Nature, 2006. 442(7105): p. 882–882.

16. Cho, S.W., et al., Targeted genome engineering in human cells with the Cas9 RNA-guided endonuclease. Nature biotechnology, 2013. 31(3): p. 230.

17. Cong, L., et al., Multiplex genome engineering using CRISPR/Cas systems. Science, 2013. 339(6121): p. 819–823.

18. Gratz, S.J., et al., Genome engineering of Drosophila with the CRISPR RNA-guided Cas9 nuclease. Genetics, 2013. 194(4): p. 1029–1035.

19. Mali, P., et al., RNA-guided human genome engineering via Cas9. Science, 2013. 339(6121): p. 823–826.

20. Hwang, W.Y., et al., Efficient genome editing in zebrafish using a CRISPR-Cas system. Nature biotechnology, 2013. 31(3): p. 227.

21. Friedland, A.E., et al., Heritable genome editing in C. elegans via a CRISPR-Cas9 system. Nature methods, 2013. 10(8): p. 741.

22. Bryant, J.M., et al., CRISPR in parasitology: not exactly cut and dried! Trends in parasitology, 2019.

23. Castelletto, M.L., S.S. Gang, and E.A. Hallem, Recent advances in functional genomics for parasitic nematodes of mammals. Journal of Experimental Biology, 2020. 223(Suppl 1).

24. Gang, S.S., et al., Targeted mutagenesis in a human-parasitic nematode. PLoS pathogens, 2017. 13(10): p. e1006675.

25. Nakayama, K.-i., et al., Screening for CRISPR/Cas9-induced mutations using a co-injection marker in the nematode Pristionchus pacificus. Development Genes and Evolution, 2020: p. 1–8.

26. Ittiprasert, W., et al., Programmed genome editing of the omega-1 ribonuclease of the blood fluke, Schistosoma mansoni. Elife, 2019. 8: p. e41337.

27. Ishino, Y., et al., Nucleotide sequence of the iap gene, responsible for alkaline phosphatase isozyme conversion in Escherichia coli, and identification of the gene product. Journal of bacteriology, 1987. 169(12): p. 5429–5433.

28. Jansen, R., et al., Identification of genes that are associated with DNA repeats in prokaryotes. Molecular microbiology, 2002. 43(6): p. 1565–1575.

29. Lander, E.S., The heroes of CRISPR. Cell, 2016. 164(1-2): p. 18–28.

30. Ledford, H., CRISPR: gene editing is just the beginning. Nature News, 2016. 531(7593): p. 156.

31. Szczelkun, M.D., et al., Direct observation of R-loop formation by single RNA-guided Cas9 and Cascade effector complexes. Proceedings of the National Academy of Sciences, 2014. 111(27): p. 9798–9803.

32. Dickinson, D.J. and B. Goldstein, CRISPR-based methods for Caenorhabditis elegans genome engineering. Genetics, 2016. 202(3): p. 885–901.

33. Qi, L.S., et al., Repurposing CRISPR as an RNA-guided platform for sequence-specific control of gene expression. Cell, 2013. 152(5): p. 1173–1183.

34. Gilbert, L.A., et al., CRISPR-mediated modular RNA-guided regulation of transcription in eukaryotes. Cell, 2013. 154(2): p. 442–451.

35. Xiao, B., et al., Epigenetic editing by CRISPR/dCas9 in Plasmodium falciparum. Proceedings of the National Academy of Sciences, 2019. 116(1): p. 255–260.

36. Walker, M.P. and S.E. Lindner, Ribozyme-mediated, multiplex CRISPR gene editing and CRISPR interference (CRISPRi) in rodent-infectious Plasmodium yoelii. Journal of Biological Chemistry, 2019. 294(24): p. 9555–9566.

37. Cámara, E., I. Lenitz, and Y. Nygård, A CRISPR activation and interference toolkit for industrial Saccharomyces cerevisiae strain KE6-12. Scientific reports, 2020. 10(1): p. 1–13.

38. Mandegar, M.A., et al., CRISPR interference efficiently induces specific and reversible gene silencing in human iPSCs. Cell stem cell, 2016. 18(4): p. 541–553.

39. Choudhary, E., et al., Gene silencing by CRISPR interference in mycobacteria. Nature communications, 2015. 6(1): p. 1–11.

40. Long, L., et al., Regulation of transcriptionally active genes via the catalytically inactive Cas9 in C. elegans and D. rerio. Cell research, 2015. 25(5): p. 638–641.

41. Li, S., et al., Enhanced protein and biochemical production using CRISPRi-based growth switches. Metabolic engineering, 2016. 38: p. 274–284.

42. Peters, J.M., et al., Enabling genetic analysis of diverse bacteria with Mobile-CRISPRi. Nature microbiology, 2019. 4(2): p. 244–250.

43. Huang, C.-H., et al., CRISPR interference (CRISPRi) for gene regulation and succinate production in cyanobacterium S. elongatus PCC 7942. Microbial cell factories, 2016. 15(1): p. 1–11.

44. Markus, B.M., E.A. Boydston, and S. Lourido, CRISPR-mediated transcriptional repression in Toxoplasma gondii. Msphere, 2021. 6(5): p. e00474–21.

45. Rock, J.M., et al., Programmable transcriptional repression in mycobacteria using an orthogonal CRISPR interference platform. Nature microbiology, 2017. 2(4): p. 1–9.

46. Whitfield, P.J. and N.A. Evans, Parthenogenesis and asexual multiplication among parasitic platyhelminths. Parasitology, 1983. 86 (Pt 4): p. 121–60.

47. Beaumier, C.M., et al., New vaccines for neglected parasitic diseases and dengue. Transl Res, 2013. 162(3): p. 144–55.

48. Lu, Z., et al., Schistosome sex matters: a deep view into gonad-specific and pairing-dependent transcriptomes reveals a complex gender interplay. Sci Rep, 2016. 6: p. 31150.

49. Wendt, G.R. and J.J. Collins, 3rd, Schistosomiasis as a disease of stem cells. Curr Opin Genet Dev, 2016. 40: p. 95–102.

50. Wang, B., J.J. Collins III, and P.A. Newmark, Functional genomic characterization of neoblast-like stem cells in larval Schistosoma mansoni. Elife, 2013. 2: p. e00768.

51. Lu, Z., et al., Schistosome sex matters: a deep view into gonad-specific and pairing-dependent transcriptomes reveals a complex gender interplay. Scientific reports, 2016. 6(1): p. 1–14.

52. Wang, B., et al., Stem cell heterogeneity drives the parasitic life cycle of Schistosoma mansoni. Elife, 2018. 7: p. e35449.

53. You, H., et al., Innovations and Advances in Schistosome Stem Cell Research. Frontiers in Immunology, 2021. 12: p. 498.

54. Collins III, J.J., et al., Adult somatic stem cells in the human parasite Schistosoma mansoni. Nature, 2013. 494(7438): p. 476–479.

55. Hahnel, S., et al., Gonad RNA-specific qRT-PCR analyses identify genes with potential functions in schistosome reproduction such as SmFz1 and SmFGFRs. Frontiers in genetics, 2014. 5: p. 170.

56. Sarfati, D.N., et al., Single-cell deconstruction of stem-cell-driven schistosome development. Trends in Parasitology, 2021.

57. Du, X., et al., Schistosoma mansoni Fibroblast Growth Factor Receptor A Orchestrates Multiple Functions in Schistosome Biology and in the Host-Parasite Interplay. Frontiers in Immunology, 2022. 13.

58. Dalton, J., et al., A method for the isolation of schistosome eggs and miracidia free of contaminating host tissues. Parasitology, 1997. 115(1): p. 29–32.

59. Du, X., et al., Gene expression in developmental stages of Schistosoma japonicum provides further insight into the importance of the Schistosome insulin-like peptide. International journal of molecular sciences, 2019. 20(7): p. 1565.

60. Oliveros, J.C., et al., Breaking-Cas—interactive design of guide RNAs for CRISPR-Cas experiments for ENSEMBL genomes. Nucleic acids research, 2016. 44(W1): p. W267–W271.

61. You, H., et al., CRISPR/Cas9-mediated genome editing of Schistosoma mansoni acetylcholinesterase. The FASEB Journal, 2021. 35(1): p. e21205.

62. Barber, R.D., et al., GAPDH as a housekeeping gene: analysis of GAPDH mRNA expression in a panel of 72 human tissues. Physiological genomics, 2005. 21(3): p. 389–395.

63. Rao, X., et al., An improvement of the 2^ (–delta delta CT) method for quantitative real-time polymerase chain reaction data analysis. Biostatistics, bioinformatics and biomathematics, 2013. 3(3): p. 71.

64. Zheng, X., et al., Soluble egg antigen from Schistosoma japonicum modulates the progression of chronic progressive experimental autoimmune encephalomyelitis via Th2-shift response. Journal of neuroimmunology, 2008. 194(1-2): p. 107–114.

65. Du, X., et al., Identification and functional characterisation of a Schistosoma japonicum insulin-like peptide. Parasites & vectors, 2017. 10(1): p. 1–12.

66. Zor, T. and Z. Selinger, Linearization of the Bradford protein assay increases its sensitivity: theoretical and experimental studies. Analytical biochemistry, 1996. 236(2): p. 302–308.

67. Bankhead, P., et al., QuPath: Open source software for digital pathology image analysis. Scientific reports, 2017. 7(1): p. 1–7.

68. Wang, T., et al., A Biomphalaria glabrata peptide that stimulates significant behaviour modifications in aquatic free-living Schistosoma mansoni miracidia. PLoS neglected tropical diseases, 2019. 13(1): p. e0006948.

69. Fogarty, C.E., et al., Comparative study of excretory–secretory proteins released by Schistosoma mansoni-resistant, susceptible and naïve Biomphalaria glabrata. Parasites & vectors, 2019. 12(1): p. 1–17.

70. Tinevez, J.-Y., et al., TrackMate: An open and extensible platform for single-particle tracking. Methods, 2017. 115: p. 80–90.

71. Wyeth, R.C., et al., Videograms: a method for repeatable unbiased quantitative behavioral analysis without scoring or tracking, in Zebrafish neurobehavioral protocols. 2011, Springer. p. 15–33.

72. Shim, M.K., et al., Caspase-3/-7-specific metabolic precursor for bioorthogonal tracking of tumor apoptosis. Scientific reports, 2017. 7(1): p. 1–15.

73. Zhu, M., et al., Inhibition of β-catenin signaling in articular chondrocytes results in articular cartilage destruction. Arthritis & Rheumatism: Official Journal of the American College of Rheumatology, 2008. 58(7): p. 2053–2064.

74. Walsh, J.G., et al., Executioner caspase-3 and caspase-7 are functionally distinct proteases. Proceedings of the National Academy of Sciences, 2008. 105(35): p. 12815–12819.

75. Eltoum, I.A., et al., Suppressive effect of interleukin-4 neutralization differs for granulomas around Schistosoma mansoni eggs injected into mice compared with those around eggs laid in infected mice. Infection and immunity, 1995. 63(7): p. 2532–2536.

76. Pearce, E.J., et al., Th2 response polarization during infection with the helminth parasite Schistosoma mansoni. Immunological reviews, 2004. 201(1): p. 117–126.

77. Zhang, P. and F. Mutapi, IgE: a key antibody in Schistosoma infection. Electron J Biol, 2006. 2: p. 11–14.

78. Gilbert, L.A., et al., Genome-scale CRISPR-mediated control of gene repression and activation. Cell, 2014. 159(3): p. 647–661.

79. Bikard, D., et al., Programmable repression and activation of bacterial gene expression using an engineered CRISPR-Cas system. Nucleic acids research, 2013. 41(15): p. 7429–7437.

80. Verkuijl, S.A. and M.G. Rots, The influence of eukaryotic chromatin state on CRISPR–Cas9 editing efficiencies. Current opinion in biotechnology, 2019. 55: p. 68–73.

81. Jurberg, A.D., et al., The embryonic development of Schistosoma mansoni eggs: proposal for a new staging system. Development genes and evolution, 2009. 219(5): p. 219.

82. Sankaranarayanan, G., et al., Large CRISPR-Cas-induced deletions in the oxamniquine resistance locus of the human parasite Schistosoma mansoni. Wellcome Open Research, 2020. 5.

83. Jamieson, B.G., Schistosoma: Biology, pathology and control. 2017: CRC Press.

84. Rinaldi, G., et al., Germline transgenesis and insertional mutagenesis in Schistosoma mansoni mediated by murine leukemia virus. PLoS pathogens, 2012. 8(7): p. e1002820.

85. Itoh, N. and D.M. Ornitz, Evolution of the Fgf and Fgfr gene families. TRENDS in Genetics, 2004. 20(11): p. 563–569.

86. Wendt, G.R. and J.J. Collins III, Schistosomiasis as a disease of stem cells. Current opinion in genetics & development, 2016. 40: p. 95–102.

87. Cheung, H.-H., X. Liu, and O.M. Rennert, Apoptosis: Reprogramming and the fate of mature cells. International Scholarly Research Notices, 2012. 2012.

88. Sarig, R. and V. Rotter, Can an iPS cell secure its genomic fidelity? Cell Death & Differentiation, 2011. 18(5): p. 743–744.

89. Costain, A.H., A.S. MacDonald, and H.H. Smits, Schistosome egg migration: mechanisms, pathogenesis and host immune responses. Frontiers in immunology, 2018: p. 3042.

90. Dutra, W.O., et al., Polarized Th2 like cells, in the absence of Th0 cells, are responsible for lymphocyte produced IL-4 in high IgE-producer schistosomiasis patients. BMC immunology, 2002. 3(1): p. 1–8.

91. Ashton, P., et al., The schistosome egg: development and secretions. Parasitology, 2001. 122(3): p. 329–338.

92. Takaki, K.K., et al., Schistosoma mansoni eggs modulate the timing of granuloma formation to promote transmission. Cell Host & Microbe, 2021. 29(1): p. 58–67. e5.

93. Yeo, N.C., et al., An enhanced CRISPR repressor for targeted mammalian gene regulation. Nature methods, 2018. 15(8): p. 611–616.

